# Neurologically altered brain activity may not look like aged brain activity: Implications for brain-age modeling and biomarker strategies

**DOI:** 10.1101/2025.04.15.648903

**Authors:** Lukas AW Gemein, Sinead Gaubert, Claire Paquet, Joseph Paillard, Sebastian C Holst, Thomas Tveitstøl, Ira RJH Haraldsen, David Hawellek, Jörg F Hipp, Denis A Engemann

## Abstract

**Background:** Brain-age gap (BAG), the difference between predicted age and chronological age, is studied as a biomarker for the natural progression of neurodegeneration. The BAG captures brain atrophy as measured with structural Magnetic Resonance Imaging (MRI). Electroencephalography (EEG) has also been explored as a functional means for estimating brain age. However, EEG studies showed mixed results for BAG including a seemingly paradoxical negative BAG, i.e. younger predicted age than chronological age, in neurological populations.

**Objectives:** This study critically examined brain age estimation from spectral EEG power as common measure brain activity in two of the largest public EEG datasets containing neurological cases alongside controls.

**Methods:** EEG recordings were analyzed from individuals with neurological conditions (n=900, TUAB data; n=417 MCI & n=311 dementia, CAU data) and controls (n=1254, TUAB data; n=459, CAU data).

**Results:** We found that age-prediction models trained on the reference population systematically under-predicted age in people with neurological conditions replicating a negative BAG for diseased brain activity. Inspection of age-related trends along the EEG power spectra revealed complex frequency-dependent alterations in neurological groups underlying the seemingly paradoxical negative BAG.

**Conclusions:** The utility of brain age as an interpretable biomarker relies on the observation from structural MRI that progressive neurodegeneration often broadly resembles accelerated aging. This assumption can be violated for functional assessments such as EEG spectral power and, potentially, different neurological and psychiatric conditions or therapeutic effects. The sign of the BAG may not meaningfully be interpreted as a deviation from normal aging.

## Introduction

Machine learning (ML) holds promise to innovate biomarker development by mapping clinical endpoints on heterogeneous biomedical imaging data^1,2^. In the context of neuroscience, the utility of ML is currently hampered by notoriously small sample sizes which are the one key limiting factor for developing ML-based prediction models^3–6^. The field has therefore turned towards exploring the potential of large public domain datasets from biobanks for training models on data from the general population to enhance the study of smaller clinical datasets^7^. In this context, applied machine-learning research has explored self-supervised learning techniques in which larger datasets are used to learn representations and features that can then be used and possibly fine-tuned for smaller datasets^8–10^. Similarly, neuroscientists have explored a number of normative approaches focusing on analysis of population-reference data which hold promise to provide biomedically interpretable outputs. This includes normative modeling^11^, brain growth charts^12,13^ and ML-derived measures of brain aging^14,15^.

Over the past decade, a substantial body of research has established brain age models based on Magnetic Resonance Imaging (MRI)^16,17^. This approach focused on estimating a function that models the relationship of chronological age to anatomical MRI features e.g. cortical and subcortical tissue statistics such as volume, surface area or thickness. These MRI features typically decline in elderly populations and neurological insults or disorders can prematurely cause brain atrophy or accelerate brain volume loss, hence increasing the apparent age of the brain^18^. As a brain-age model calibrated on the general population is applied to neurodegenerative populations, systematic overestimation can occur as the brain of a patient may appear “older” in comparison to the population reference^19,20^. This is the core idea captured by the brain-age gap (BAG) which is computed by subtracting the chronological age of a person from the MRI-predicted age^21^, such that positive values represent a higher biological than chronological age of the brain. In patients suffering from different neurological conditions a positive BAG has been consistently reported^22–25^ and a higher BAG even predicted future mortality^26^, highlighting the potential of the BAG as a prognostic biomarker. Such metrics could be useful as stratifiers or endpoints in clinical research including interventional studies targeting neurological conditions. A model-based brain-age approach leverages large datasets from the general population and applies ensuing models to research questions with much smaller groups. These models may better summarize baseline brain health or track disease progression in terms of progressive cerebral atrophy.

A more recent line of research has explored the utility of brain-activity measures for brain-age modeling^27^. Studies focusing on functional MRI (fMRI) have pointed out that functional-connectivity measures contribute complementary information over anatomical features which can lead to improved clinical characterization^28^. More recently, electroencephalography (EEG) and magnetoencephalography (MEG) have also been explored as a modality for studying brain aging with ML. M/EEG measures large-scale synchrony of cortical neurons across multiple temporal scales from milliseconds to hours^29,30^, hence, providing a unique view on brain function and health. Multimodal studies have found that MEG contributes unique information on age compared to fMRI and MRI^31,32^ and several publications have demonstrated successful age-prediction from EEG signals^33–37^ with performance scores somewhat inferior to those obtained using anatomical MRI while adding complementary physiological information.

These results may suggest that EEG could serve as an attractive solution for applying brain-age models in real-world settings, as the ease of access to EEG can enable its use in a wide range of societal and medical situations. In fact, besides laboratory-based resting-state EEG, age-prediction has been explored with sleep EEG^33,37^, EEG from anesthetic monitoring^38^ and at-home recordings during sleep or meditation EEG^37^. However, the implied clinical significance has rarely been systematically explored and many studies do not present separate training on healthy reference populations and testing on clinical populations^35,39^, hindering clinical interpretation in the context of brain age.

Supportive evidence for the utility of EEG based brain-age was provided by a line of research focusing on laboratory-based sleep EEG with around 5-8 years of mean absolute error in large baseline datasets^33,40–42^, which falls within the range of brain-age benchmarks for wakeful resting EEG and MEG^34^. Applications of these models to cohorts of dementia patients showed a positive BAG – in line with an interpretation as a measure of brain aging – and reported that brain age was increasingly elevated in dementia patients^40,42^. Another related study found elevated sleep-EEG brain age as a risk factor of mortality^41^.

Mixed evidence was put forward by another line of work reporting seemingly paradoxical effects as patients appeared to show a “younger brain age”, suggesting that brain activity changes in certain neurological and anaesthetized conditions may oppose those changes seen with aging^38,43^. Using a deep convolutional neural network applied to raw EEG signals, Gemein et al.^43^ found a negative BAG, suggesting “younger” brain age, in patients with EEG anomalies in the Temple University Hospital Abnormal EEG corpus (TUAB)^44,45^. Likewise, using Riemannian techniques capturing spectral power, Sabbagh et al.^38^ found that among older patients undergoing propofol anesthesia, those who showed anesthetic anomalies such as iso-electric burst suppression^46,47^ showed lower brain-age compared to those who did not – implying an induced negative, “younger”, BAG through anesthesia.

Numerous studies have documented abnormal changes in brain activity in e.g. the course of Alzheimer’s disease related to amyloid and tau pathology^48–53^. Complex pathological alterations in local circuits and brain networks lead to reorganization of brain activity implying substantial differences in aging-related patterns between neurological populations. This raises the question whether the normative brain-age concept related to the earlier onset of cerebral atrophy readily generalizes to measures of brain function, such as the EEG, which may be altered in complex ways as compared to healthy aging.

We used two large public datasets and studied predictions from brain age models trained on EEG power spectra from healthy reference populations when applied to neurological populations. We subsequently explored the relationship between age and signal power across the EEG power spectrum. This allowed us to develop the hypothesis that complex age-trends of functional biomarkers in neurological populations facilitate brain-age predictions of unexpected directionality. Our findings may inform alternative strategies and potential solutions for EEG-based ML and its clinical applications.

## Results

### EEG-based brain-age models trained on healthy controls predicted substantially younger brain age for neurological individuals

We constructed age-prediction models on EEGs using two large public datasets containing neurological EEG data alongside controls, the Temple University Hospital Abnormal EEG corpus (TUAB, n = 2154)^44,45^ and the Chung-Ang University Hospital EEG (CAU, n = 1187)^54^ dataset. The TUAB dataset contained heterogenous archival data from patients seeking neurological counseling with precise diagnoses unknown for privacy reasons, which includes an important amount of epilepsy-related anomalies. The subset of TUAB containing anomalies is commonly referred to as “pathological” in the literature. The CAU dataset contains patients diagnosed with mild cognitive impairment and various forms of dementia. Here we focused on Alzheimer’s dementia cases (the dominating subcategory). As the age-range in CAU was limited, we pooled the control populations across both datasets to obtain a wider age distribution and use as many training samples as possible. Applying a regularized linear prediction model (ridge regression) we obtained age predictions with reasonable calibration (R^2^=0.61; Figure 1a), in line with state-of-the art benchmarks for MEG and EEG.^34,43,55^ We then investigated the ensuing brain age gap (BAG) for neurological recordings (Figure 1b). We observed that, on average, the EEG-predicted age was between -10 and -6 years for neurological EEG whereas control groups showed predictions centered around zero (Figure 1c, left panel). The point clouds (Figure 1a) showed a wider spread for neurological groups, suggesting lower prediction performance that may reflect increased biomedical heterogeneity.

**Figure 1.**
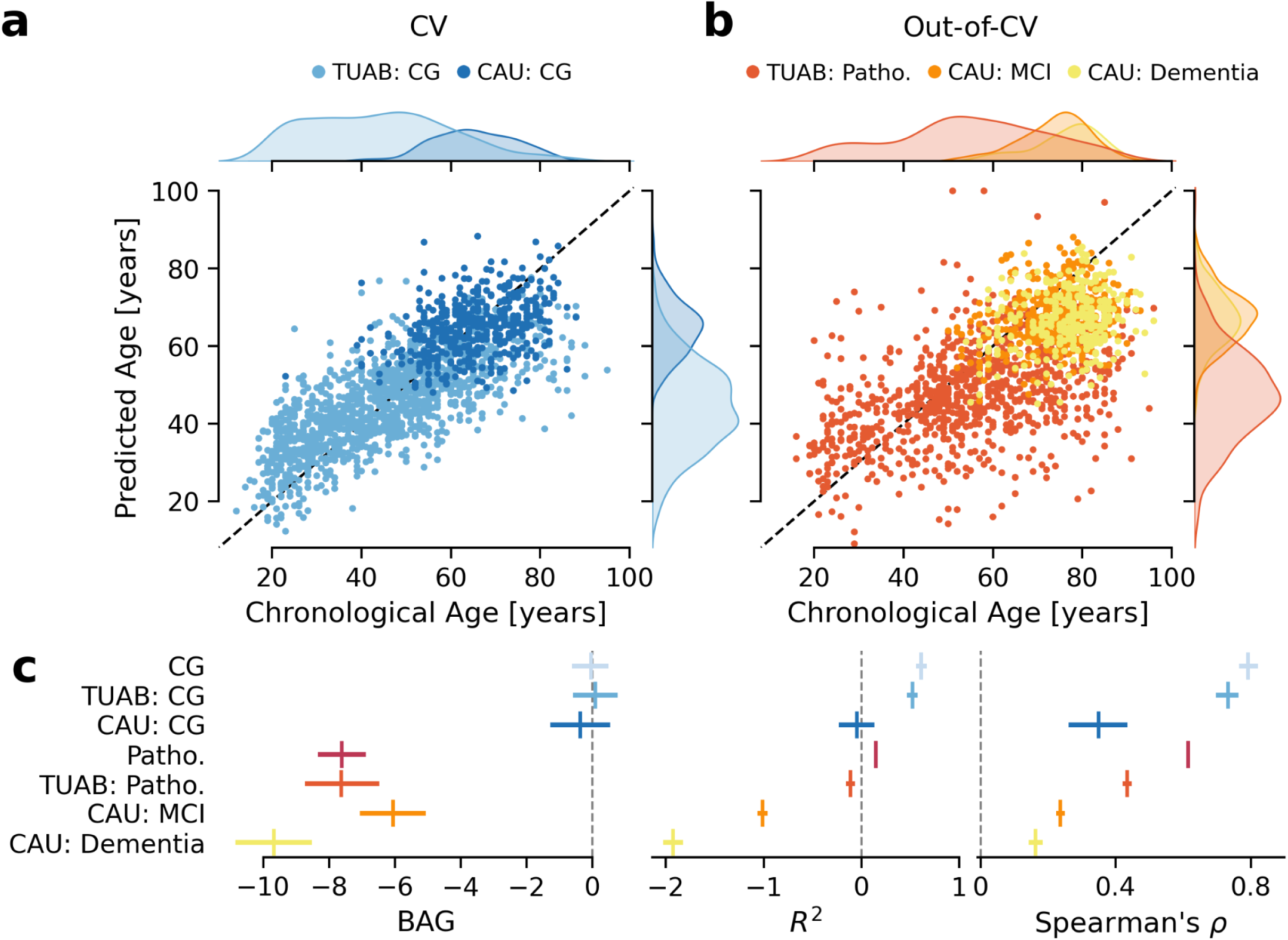
Brain-age models derived from EEG power spectra of controls predicted younger age in neurological cases. (**a**) Chronological versus predicted age for the joint control groups of the Temple University Hospital Abnormal (TUAB) and Chung-Ang University Hospital (CAU) datasets in a 10-fold cross-validation. Both datasets align along the diagonal, indicating accurate predictions, with a slight underestimation trend in older individuals. (**b**) Chronological versus predicted age for the neurological groups of TUAB and CAU. Compared to the control group (CG), panel (a), the fit was poorer, with a more pronounced underestimation. (**c**) BAG scores for individual conditions and overall control and neurological groups. Control groups have BAGs centered around zero, indicating well-calibrated models. Neurological groups showed consistently negative BAGs, with CAU/dementia appearing the "youngest" on average. Despite negative *R*^2^ scores for CAU/dementia and CAU/MCI, samples were still meaningfully ranked across conditions as indicated by Spearman’s correlation ρ.

To rule out technical errors, we examined prediction performance using the R^2^ performance metrics that are sensitive to translation and scaling errors and Spearman’s rank correlation that is robust to these errors and captures non-linear correlations (Figure 1c). In line with negative brain age (Figure 1a), constituting a translation error, the R^2^ scores below zero indicated absence of model fit (Figure 1c, mid panel). On the other hand, Spearman’s rank correlations reveal positive association between brain-age predictions and the actual age of the patients (Panel 2c, right). This suggests that the models learned, to some extent, systematic patterns from the data that generalize to the neurological population despite a global negative shift in BAG scores and that negative BAG scores did not occur due to global prediction failure, i.e., random guessing.

A limitation of our analysis arises from the substantially higher age of neurological groups in the TUAB and CAU datasets (Figure 1a-b, marginal histograms), rendering age a strong differentiator between the groups. Unfortunately, no neurological scores or other clinical variables are available for the two datasets that could enable further contextualizing the BAG scores. To gauge the clinical utility of the BAG scores for distinguishing between groups while controlling for age, we constructed statistical logistic regression models, additively combining age and brain-age for separating cases and controls (Figure 2, for details cf. *Statistical Analyses* in *Methods*). To account for potential nonlinear effects, we also included age^2^ as was done in previous work.^7,31^

**Figure 2.**
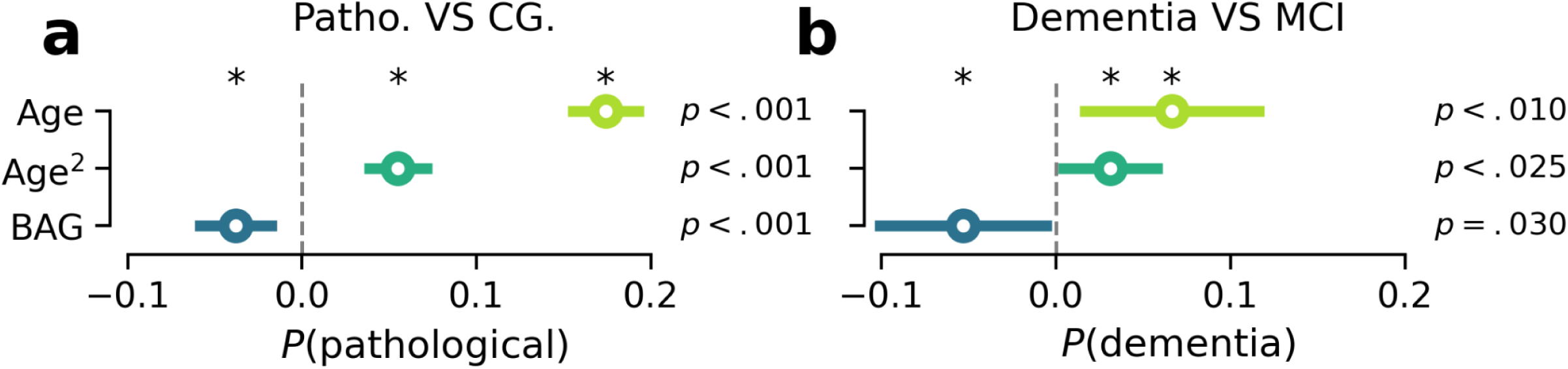
EEG-derived BAG captured information complementary to age but with unexpected directionality. Logistic regression analysis of chronological age and brain-age gap on the y axis with marginal effects (change in predicted probability) on the x axis for (**a**) differentiating CG from neurologically abnormal groups and (**b**) MCI from dementia. All variables provided significant independent information. Increased age and age^2^, and, conditional on the age terms, a smaller BAG (suggesting a "younger" brain age) was associated with a higher probability of EEG anomalies.

We found that the BAG provided additive information beyond linear and nonlinear functions of age for differentiating EEG with anomalies from controls in the TUAB data (Figure 2a) and dementia from MCI cases in CAU data (Figure 2b). However, consistently, across the age range (conditional on age and age^2^), negative BAG (“younger” looking EEG) was associated with a higher predicted probability of EEG anomalies (Figure 2a) or neurological cases (Figure 2b). These two examples demonstrate that EEG-derived brain-age contains certain statistical signals. However, these results highlight the risk of misinterpretations as one would not expect that neurological conditions lead to signatures of brain-activity resembling younger healthy people.

Taken together, these results replicate recent literature observing unexpected directionality in brain-age predictions using EEG. We observed a negative BAG in populations for which one assumes lower brain health and advanced biological age^38,43^.

### Neurological EEG shows complex increases and decreases in brain activity

To better understand the negative (younger) BAG in elderly neurological populations, we inspected age-related trends in the data used as input to the age-prediction models. It is in this context worthwhile to note that MEG and EEG studies investigating the brain signals driving age prediction pointed out that information used by brain-age models was wide-spread across the entire power spectrum and not concentrated on specific frequency windows.^31,37,38,56,57^ This means that if age-related trends for key features are modified or even inverted in specific frequencies, a model trained on healthy references, paying attention to all frequencies, will be affected and may fail to provide an accurate estimate of the relationship between EEG and age.

We therefore investigated differences in power spectral densities (PSD) between the case and the control groups (Figure 3). In both datasets (TUAB and CAU) and across all conditions (EEG anomalies, MCI, dementia), we observed simultaneous decrease and increase in EEG power depending on the frequency when comparing cases to controls (Figure 3a). Specifically, for neurological groups we found higher power in low frequencies (up to ∼8 Hz) and lower power in medium frequencies (∼8-32 Hz) as compared to controls (Figure 3a, right panel).

**Figure 3.**
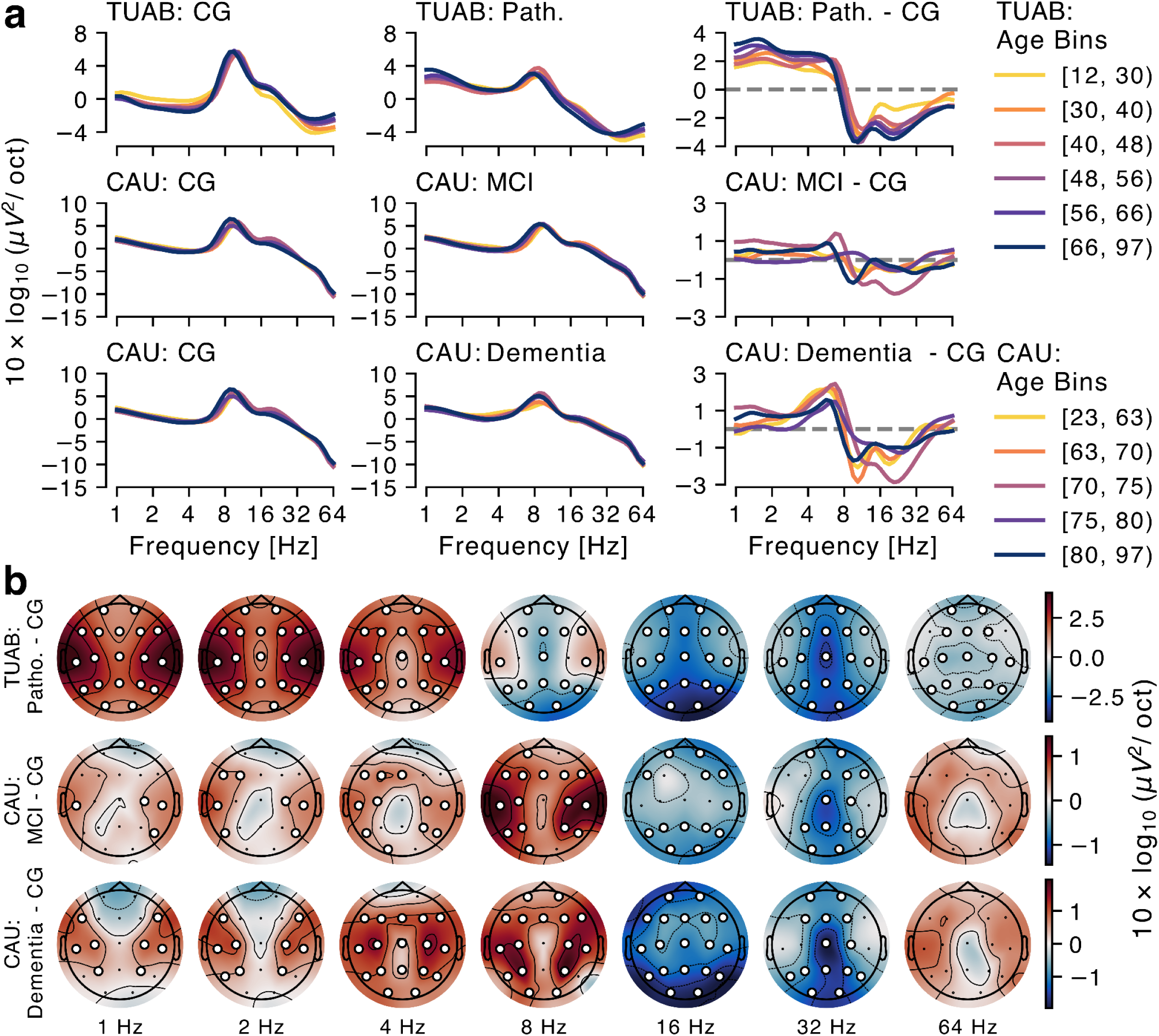
EEG power spectra are differently modulated across age for neurological cases and controls. (**a**) Average log EEG power by age groups for cases with and without pathological anomalies and comparisons in TUAB, CAU/MCI, and CAU/dementia. Neurological recordings showed higher power below 8 Hz and lower power between 8 Hz and 32 Hz across all age ranges, most prominently in TUAB, followed by CAU/dementia and CAU/MCI. (**b**) Topographic maps of power differences between cases and controls in TUAB, CAU/MCI, and CAU/dementia at octave frequencies. White markers show significant differences (p < 0.05) between the groups, TFCE permutation test for a T-statistic. In TUAB and CAU, topographic maps revealed significantly higher power below 8 Hz and significantly lower power between 8 Hz and 32 Hz for pathological recordings. This effect was globally extended across frequencies in TUAB and more spectrally localized in CAU/dementia. Applying brain-age methodology, higher power in afflicted individuals would be unexpected as power naturally declines with age and neurological disorders typically increase the rate of decline.

To formalize these observations statistically and to include systematic tests for the relationship between EEG and age, we designed a repeated-measures 3-way ANOVA model of mean EEG power with the factors group, frequency range (low: <16Hz, high: >=16Hz), and age and all interactions thereof (Table 1). For both datasets, multiple significant interactions were obtained, pointing at a complex data scenario: Conditional on all other model terms, average power differed between neurological groups and high frequencies. This is a directly related consequence of our visually guided choice of the 16Hz cut-off and octave normalization removing 1/f trends. Importantly, in both datasets, we observed independent two-way interactions suggesting non-linear changes of power with age in higher frequencies and for neurological groups. Importantly, for TUAB, the three-way interaction term revealed that the direction of correlation of power with age was significantly altered in high versus low frequencies depending on the group.

**Table 1.** Repeated measures ANOVA of log power by age, group (neurological VS control) and frequency (high VS low), separately by dataset.

| Source | TUAB |  |  |  |  | CAUEEG |  |  |  |  |
| --- | --- | --- | --- | --- | --- | --- | --- | --- | --- | --- |
|  | DF | SSQ | MSQ | F | p | DF | SSQ | MSQ | F | p |
| **Group** | 1 | 314.0 | 313.99 | 667.832 | <<br>**0.001**<br>*** | 1 | 3.80 | 3.802 | 14.021 | <<br>**0.001**<br>*** |
| **Age** | 1 | 2.6 | 2.58 | 5.489 | **0.019**<br>* | 1 | 3.32 | 3.322 | 12.253 | <<br>**0.001**<br>*** |
| **Group X Age** | 1 | 15.3 | 15.30 | 32.538 | <<br>**0.001**<br>*** | 1 | 0.20 | 0.201 | 0.741 | 0.390 |
| **Residuals (between)** | 2150 | 1010.9 | 0.47 |  |  | 763 | 206.89 | 0.271 |  |  |
| **frequency** | 1 | 618 | 618 | 123.093 | <<br>**0.001**<br>*** | 1 | 546.3 | 546.3 | 188.819 | <<br>**0.001**<br>*** |
| **Group X Frequency** | 1 | 3351 | 3351 | 667.832 | <<br>**0.001**<br>*** | 1 | 40.6 | 40.6 | 14.021 | <<br>**0.001**<br>*** |
| **Frequency X Age** | 1 | 28 | 28 | 5.489 | **0.019**<br>* | 1 | 35.5 | 35.5 | 12.253 | <<br>**0.001**<br>*** |
| **Group X Frequency X Age** | 1 | 163 | 163 | 32.538 | <<br>**0.001**<br>*** | 1 | 2.1 | 2.1 | 0.741 | 0.390 |
| **Residuals (within)** | 2150 | 10787 | 5 |  |  | 763 | 2207.7 | 2.9 |  |  |

*Note.* Significance levels: \*\*\*: $p < 0.001$ , \*\*: $p < 0.01$ , \*: $p < 0.05$ , . : $p < 0.1$ ; DF: degrees of freedom, SSQ: sum of squares, MSQ: mean square

Against this background, it is worthwhile inspecting topographic maps which revealed statistically significant differences (𝑝 < 0. 05) on a large coherent set of electrodes across a wide range of frequencies (Figure 3b). At around 16 Hz, we noticed an inversion of significant activation differences (Figure 3b). In line with the interaction effects (Table 1), it can be seen that power differences between cases and controls, across age groups, could show opposite trends, depending on the frequencies (Figure 3a, right subpanels). For example, for TUAB, as indexed by color codes (Figure 3a), the oldest group (66-97 years) showed smaller positive (below 16 Hz) and smaller negative (∼16-32 Hz) differences, whereas younger groups showed larger positive differences (below 16 Hz) and larger negative differences (∼16-32 Hz).

These analyses demonstrate that neurological anomalies were associated with diverse and complex deviations from the EEG signals of age-matched controls along the frequency spectrum. The observed signature of higher power in low frequencies and lower power in high frequencies resembles the commonly studied cortical slowing phenotype characteristic of clinical stages of Alzheimer’s Dementia^58–60^ but was also observed for epilepsy.^61^ Taken together, this suggests that EEG power spectra from neurological patients do not simply look like prematurely aged – “old” – EEG power spectra from healthy reference populations.

### Explaining the paradoxical BAG with differences in cross-sectional EEG trends for cases versus controls

We have two main findings so far: We observed unexpected directions of association between EEG-based brain-age predictions in neurological cohorts. And we observed complex differences in EEG power spectra between neurological cases and healthy controls. Confronting these findings with another provides us with a foundation for theoretically explaining factors causing systematic miscalibration in brain-age models. The core of our emerging hypothesis for brain-age models that can give counter-intuitive directions of prediction (i.e. younger brain age in patients with neurodegenerative conditions) is that neurodegeneration leads to cross-sectional aging trends in structural MRI that parallel aging trends seen during aging in controls, however, earlier and faster^7,14,15,18,25^. In other words, for MRI, changes due to disease and aging are similar and differ mostly in their timing and their longitudinal slope (Figure 4a). Given the rich and complex brain signals captured by EEG, neither the characteristics of the changes nor their time constants have to be necessarily similar between disease and normal aging (Figure 4b). As a result, differences between diseased brain activity and aged brain activity can occur in multiple directions at the same time. As the brain-age predictions are based on models trained with healthy controls, they can be viewed as a mapping to the nearest neighbour in the healthy reference population. An unexpected directionality of BAG scores can occur if patients’ EEG data violate the pattern with which age is driving EEG changes and, for example, exhibit higher levels of brain activity that would be otherwise typical of younger healthy controls. The EEG signature observed in Figure 3 for neurological groups relative to healthy controls^53,59,60,62–64^ could represent such a scenario.

**Figure 4.**
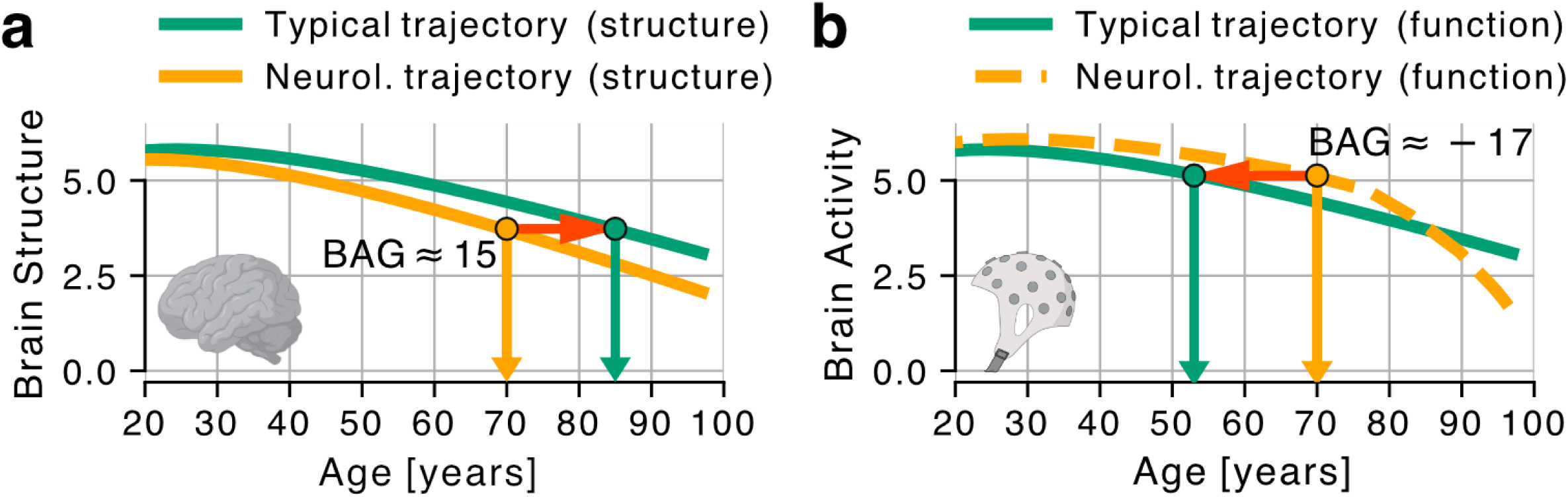
Emerging hypothetical explanation for brain-age trends with unexpected directionality. (**a**) In the first scenario, the pathological biomarker trajectory decays earlier compared to the typical population but follows a similar, mainly translated trajectory. A brain-age model trained on normative data overestimates the age of a 70-year-old individual (positive BAG), common with MRI features reflecting brain atrophy. (**b**) Here, the pathological biomarker trajectory initially deviates positively then negatively from the typical trajectory. The model underestimates the age of a 70-year-old individual (negative BAG), a scenario possible with EEG power measures that increase abnormally before declining, as seen in progressive MCI^71^. This can result in sign ambiguity in brain-age models, complicating the interpretation of candidate measures of EEG-derived brain age.

But other scenarios could lead to similar effects, e.g., when not controlling for medication that may affect spectral power, such as allosteric GABA modulators ^65–67^ (e.g. benzodiazepines) as well as differences in baseline physical activity^68^. As a consequence, it is important to appreciate that neurological diseases, even if involving “pathological aging” are not necessarily the same as earlier and faster aging. Neurological conditions such as Alzheimer’s disease or epilepsies are caused by a complex interplay of genetics and life history, leading to physiological processes that do not occur equally in unaffected people.^69,70^ One, therefore, needs to be careful which assumptions remain valid from a model trained on healthy controls when applied to specific neurological populations. This is more than merely an empirical question as it becomes a scientific question given the cumulative progress in deciphering the altered physiological processes and mechanisms involved in major neurological conditions.

## Discussion

In this work, we explored potential pitfalls of EEG-based brain age models, revisiting counterintuitive findings according to which neurological conditions or anomalies were associated with lower rather than higher brain-predicted age^43,72^. Our literature review, empirical benchmarks and conceptual analysis argued that under neurological conditions, progressive changes in EEG can be different from those associated with normal aging in complex ways. This impedes a simple generalization of the MRI-based brain-age framework to EEG.

MRI-based brain-age makes the assumption that the brain changes through neurodegeneration are similar in their pattern but earlier and faster versions of the brain changes with aging^7,14,15,18,2518^. Across large and diverse clinical cohorts including more than 3,000 EEGs, we demonstrated that age-related changes in EEG power showed complex signal alterations due to different neurological conditions with simultaneous increases and decreases in brain activity compared to age-matched controls (Figure 3). In particular, the analyses showed that unlike for structural MRI showing earlier atrophy with a similar pattern to an older brain, a neurological EEG does not necessarily look like an EEG of an older brain. Using the same power spectral data, we found that brain-age models trained on the control groups showed systematically negative associations between brain-age and neurological cohort, suggesting that neurological anomalies or even neurodegeneration would paradoxically go along with a “younger” brain age. These observations stimulated our conceptual analysis aiming to explain the unexpected directionality in associations between brain-age and objective brain-health in terms of complex altered aging trends in EEG that can be substantially different in disease as compared to health. Together, our results illustrate the hidden complexity of brain-age modeling with functional data and raise the question how to best support the endeavor of exploring neurological biomarkers with rich brain-activity measures using machine learning.

One important motivation for the brain-age methodology has been the idea of enhancing the analysis of smaller datasets of limited size in the context of specific applications (e.g. a clinical trial) by applying machine learning models on population-level large-scale. Anatomical brain age has been explored for this end^7,25^ as it has the advantage of providing an intuitive summary of structural changes and it may be an effective way to augment datasets where normal aging and pathology share common effects. It is important to note that emerging data from the pharmacological context suggests that the effects of study drugs may introduce similarly paradoxical effects for structural MRI as well. For example, monoclonal antibody therapies for Alzheimer’s disease were shown to induce an unexpected additional rate of brain atrophy^73,74^ and treatment with antisense oligonucleotides can be associated with an expansion of the size of the ventricular system^75^. These changes occur even in the face of an established clinical benefit in the case of the antibody treatments. Accelerated brain atrophy signals associated with a clinical benefit suggest a complex difference between brain-age predictions and changes in the objective brain-health due to treatment for structural MRI. Thus, a new conceptual framework for describing the link between brain-age and brain-health may not only be needed for functional measures but also unexpected treatment related changes.

As the number of larger datasets for model training keeps growing also for EEG, it becomes increasingly practical to train models directly on targets of interest (e.g. diagnosis) rather than constructing a proxy via brain age. Our results on classification of CG vs. Pathological in TUAB or CG vs. MCI vs. dementia in CAU add to the growing body of literature^56,76–79^ highlighting the capability of EEG for directly modeling diverse targets of interest by capturing complex and fine-grained signal alterations in frequency and space. It is conceivable that exploring the wealth of existing machine learning models and methods for domain adaptation and transfer learning^80–84^ could lead to scenarios with reasonable brain age applications. But against the background of the methodological issues exposed in this work, the indirect brain age modeling approach – applied to EEG – may not deliver on the promise of a simple and interpretable clinical tool that tells how “old the brain looks”^16,25,85^. Meanwhile, the brain age approach could still be useful in conjunction with other data as a proxy measure by adding statistical information to smaller clinical datasets even if falling short of providing an actionable biomarker^7^. A recent study systematically investigated the advantage of MRI-derived brain-age models versus direct models of Alzheimer’s Dementia and did not report any advantages of the indirect brain-age-based approach over direct prediction of dementia-specific health outcomes.^86^ Future work will be needed to further consolidate these results across signal modalities and disease areas.

It is important to note that some applications of EEG-based brain-age showed expected results with higher brain-age in dementia patients.^33,40–42^ These disparities grant the opportunity for hypothetical reflections on potential underlying mechanisms of calibration, i.e., accurate generalization performance, in brain-age predictions. Fundamentally, calibration suggests that the prediction functions are of a similar shape and only translated, e.g. showing similar aging trends in key features (Figure 4a). Successful calibration observed in studies involving sleep data ^33,40,41^ might, thus, imply that some of the sleep characteristics e.g. amount of deep sleep, slow wave activity, sleep spindles and sleep fragmentation may lie on parallel trajectories in healthy and pathological aging and that those features were picked up by the prediction models. This would be consistent with recent observations detailing positive associations between sleep impairments and markers of neurodegeneration, including mild cognitive impairment^87^, alongside tau and beta amyloid levels in the cerebrospinal fluid ^88,89^. Interestingly, even a single night without sleep can enhance cerebrospinal fluid markers of beta amyloid and tau by more than 35% and translates into an MRI based brain age increase of 1 to 2 years.^90,91^ On the other hand, it is possible that EEG-derived brain-age modeling based on anatomy-informed source imaging extracts a trajectory that is constrained by the MRI trajectories, hence, inducing calibration. In a recent study reporting calibrated brain-age predictions from EEG, neural interaction features related to mutual information metrics estimating synergy vs. redundancy between brain regions conditional on all other interactions were used as model input.^42^ The characteristic feature was thus by construction independent from overall brain-activity differences and instead, potentially, isolated changes in the spatial structure of the data, related to the integrity of long-range neuronal connections. The focus on spectral power and simple regularized linear prediction models in this study can be seen as a limitation of our study. But on the other hand, miscalibration has even been observed with complex deep learning architectures that can learn sophisticated features from raw EEG traces.^43^ Regardless, none of this affects the principal point developed in this work that calibration of brain-age models is not self-evident and requires an explicit strategy or strong empirical sensitivity analysis. And then, not even well calibrated models are guaranteed to be useful.^86^

Going one step further, we would argue that the intrinsic richness and versatility of EEG at detecting functional anomalies longs for intensified modeling both in neurological individuals and control groups beyond ML-based classification. For example, normative modeling^11,12,92^ aims to create statistical descriptors of normal variations and can therefore be used to identify individuals that deviate therefrom. Related brain growth charts aim to describe development of the brain throughout the human lifespan^13^ and event-based modeling aims to identify the sequence and timing of progression to improve understanding of the disease and to identify possible points for intervention.^64,93^ Therefore, the identification and definition of informative EEG biomarkers would be greatly facilitated by carefully assessing disease trajectories and their divergence to normative baselines^92^ by inferring disease stages and progression of severity.^64,93–96^

This leads to a final consideration. Despite the constantly growing public availability of sources of large amounts of training data, the field continues to face challenges of limited data availability. This has even affected our current work as limitation as no detailed diagnoses were made available for TUAB and diagnoses for Alzheimer’s disease were purely clinical for the CAU data. However, combining multiple public domain datasets, as we performed this work, is an important first step. A fundamental implication of this study is that large samples from healthy reference populations are not sufficient. In addition, to perform trajectory modeling or direct classification, large datasets describing the clinical population and problem of interest are required. Therefore, the significance of ML for EEG comes down to the availability of trusted data – the *right data* and not only the *big data*. For one, there might be hidden potential in the process of examination, i.e. increasing data quality by limiting the influence of adverse behavior such as physical exercise^97^, and repeated assessment to cover intra-subject variability^72^. For another, targeted studies of specific diseases and disorders with longitudinal assessments^98^ are key to advancing the acceptance of EEG in clinical practice and as a trusted source for deriving robust biomarkers. To make a difference, we believe that the community must come together and focus on overcoming the impediments for collecting large-scale neurological data with high-quality EEG.

## Methods

### Data

In this work, we used recordings of the Temple University Hospital Abnormal EEG (TUAB) (CG: n=1254, Pathological: n=900)^44,45^ and the Chung-Ang University Hospital EEG (CAU) (CG: n=459, MCI: n=417, and dementia: n=311)^54^ corpora. The age range of CG in TUAB is 12-95 years while it is 16-96 years for the so-called “pathological” group. As precise diagnoses are not available for this dataset and the recordings from that latter group contain heterogeneous anomalies stated by trained medical observers, the designation “pathological” is commonly used in the literature using the TUAB dataset. The age range of CG in CAU is 23-88 years, 50-93 years in MCI, and 54-96 years in dementia. We refer to individuals with normally/healthy/non-pathologically labeled EEG recordings as control group (CG) and to individuals with a pathologically/MCI/dementia labeled recording as a neurological group. It has to be noted, though, that all recordings took place in a clinical setting, i.e. were recorded from individuals that came to a hospital and where staff decided to conduct an EEG examination. Furthermore, within the context of this work, the terms recordings and individuals are used interchangeably, as only a single recording of each individual was included.

### Machine learning pipeline

#### Preprocessing

The datasets were harmonized as closely as possible in terms of the individual’s state (resting), sampling frequency (250 Hz), referencing (common average), channel number (19) and order (alphabetically). We applied thorough preprocessing established in other works^99,100^ including bad segment rejection and epoch repair through autoreject^101,102^, removal of artifact sources via ICA^103^, accounting of inter-individual variability of global amplitude levels with the global scale factor^104,105^.

#### Feature extraction and model construction

We computed covariance matrices based on EEG signals filtered with log-linearly spaced Morlet wavelets^99,106^ which we transformed to log power features. We used ridge regression with hyperparameter tuning via generalized cross-validation which has proven highly effective and simple in concert with power-spectral features^34,35,99^. We applied the same model configuration as in previous work^99^.

### Model evaluation

We conducted 10-fold cross-validation on the joint control groups of TUAB and CAU. Therefore, data was shuffled and stratified with respect to the dataset of origin. When building the joint model, to account for distribution differences between the datasets and groups, we included a dataset indicator to fit a dataset-specific intercept term. During every fold, data (log power values and site indicator) as well as targets (age) were scaled to zero mean and unit variance with respect to the training set. In addition to the control groups, we evaluated the linear model (RidgeCV) on all neurological groups of both datasets.

### Brain age

We computed BAG as predicted - chronological age to obtain intuitively interpretable measures of underestimation (negative sign) or overestimation (positive sign) of chronological age. We computed 𝑅 scores to quantify the proportion of variance explained by our linear regression model. Additionally, we computed Spearman’s rank correlation coefficients to assess the strength and direction of the monotonic relationship between the predictor and target variables.

### Statistical analysis

In this study, we employed a logistic regression model to investigate the relationship between pathology status (control vs. pathological in TUAB and MCI vs. dementia in CAU) and predictors chronological 𝑎𝑔𝑒, ^2^ and brain age gap (BAG). The logistic regression model was motivated by previous works’ methods for analyzing the complementarity of age versus brain age^7,31^ and was specified as follows:

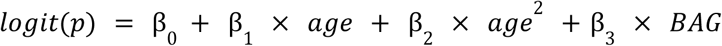

where 𝑝 denotes the probability of a specific outcome, β_0_ is the intercept, and β_1_, β_2_ and β_3_ are the coefficients for chronological 𝑎𝑔𝑒, age^2^ and brain age gap, respectively.

To interpret the impact of each predictor on the probability of the pathology status, we assessed the marginal effects ^107,108^. Marginal effects provide insights into how changes in the predictors are associated with changes in the predicted probability 𝑝 which is of greater interest here than the latent linear predictor. The coefficients β_1_, β_2_ and β_3_ capture the linear effects of the predictor variables on the log-odds of the outcome. A positive coefficient indicates that with increasing value, the log-odds of the outcome increase, suggesting a higher probability of the outcome with increasing coefficient. Conversely, a negative coefficient would indicate a decrease in the log-odds with increasing coefficient. We considered results significant at a significance level of 0.05.

We compared spatial power distributions of neurological groups with respect to their control groups with spatial-spectral permutation tests adapted from MNE^109^, essentially performing an independent sample t-test and threshold-free cluster enhancement^110^ with 10000 permutations. We visualized the resulting topographic maps with markers at significant electrode locations at a significance level of 0.05.

To explore age-related trends in the power spectra, we performed a repeated measures ANOVA for the TUAB and CAU datasets expanding all model terms for the following model formula syntax:

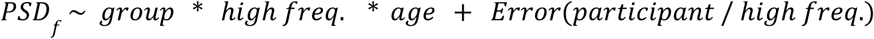

*High frequency* was defined as frequencies above 16Hz and motivated by visual exploration revealing a reversal of the sign of power differences between groups (Figure 3). This categorical term was necessary to express complex reversal patterns that cannot be described in terms of mere product interactions between continuous frequency and age – at least not without use of polynomial terms or spline features leading to more complex models that are harder to interpret. Intuitively, interaction terms involving this categorical *high frequency* variable allows fitting independent linear slopes for high versus low frequencies.

Ethics

### TUAB

We cite the statement from the original data release publication^44^: “All work was performed in accordance with the Declaration of Helsinki and with the full approval of the Temple University Institutional Review Board (IRB). All personnel in contact with privileged patient information were fully trained on patient privacy and were certified by the Temple IRB”.

### CAU

An official research proposal was submitted and accepted by the Chung-Ang University Hospital ethics committee.

### Software

All code was implemented in Python 3.11.

Main libraries used were Matplotlib 3.9.0, MEEGLET 0.0.1rc7, MNE-Python 1.7.0, NumPy 2.0.0, Pandas 2.2.2, Seaborn 0.13.2, scikit-learn 1.5.0, and statsmodels 0.14.2.

Illustrations in Figure 4 were partially created with biorender.com

## Code and Data Availability

Both the TUAB and the CAU dataset are publicly available upon registration (https://isip.piconepress.com/projects/nedc/html/tuh_eeg/#c_tuab, https://github.com/ipis-mjkim/caueeg-dataset).

## Declaration of Interests

Lukas AW Gemein, Joseph Paillard, Sebastian Camilo Holst, David Hawellek, Jörg F Hipp and Denis A Engemann have been full-time employees of F. Hoffmann-La Roche Ltd.

## Contributions

**Conceptualization:** LAWG, SG, JFH, DAE

**Data Curation:** LAWG, JP, DAE

**Formal Analysis:** LAWG, DAE

**Funding Acquisition:** DAE

**Investigation:** LAWG

**Methodology:** LAWG, DAE

**Project Administration:** DAE

**Resources:** JH, DAE

**Software:** LAWG, JH, DAE

**Supervision:** DAE

**Validation:** LAWG, SG, CP, JP, JH, DAE

**Visualization:** LAWG, DAE

**Writing - Original Draft:** LAWG, DAE

**Writing - Review & Editing:** LAWG, SG, CP, JP, TT, IRJHH, DH, JH, DAE

## Notes

https://isip.piconepress.com/projects/nedc/html/tuh_eeg/#c_tuab

https://github.com/ipis-mjkim/caueeg-dataset

### Competing Interest Statement

Lukas AW Gemein, Joseph Paillard, Sebastian Camilo Holst, David Hawellek, Joerg F Hipp and Denis A Engemann have been full-time employees of F. Hoffmann-La Roche Ltd.

